# *NGMASTER: in Silico* Multi-Antigen Sequence Typing for *Neisseria gonorrhoeae*

**DOI:** 10.1101/057760

**Authors:** Jason C Kwong, Anders Gonçalves da Silva, Kristin Dyet, Deborah A Williamson, Timothy P Stinear, Benjamin P Howden, Torsten Seemann

## Abstract

Whole-genome sequencing (WGS) provides the highest resolution analysis for comparison of bacterial isolates in public health microbiology. However, although increasingly being used routinely for some pathogens such as *Listeria monocytogenes* and *Salmonella enterica*, the use of WGS is still limited for other organisms, such as *Neisseria gonorrhoeae*. Multi-antigen sequence typing (NG-MAST) is the most widely performed typing method for epidemiologic surveillance of gonorrhoea. Here, we present *NGMASTER* – a command-line software tool for performing *in silico* NG-MAST on assembled genome data. *NGMASTER* rapidly and accurately determined the NG-MAST of 630 assembled genomes, facilitating comparisons between WGS and previously published gonorrhoea epidemiological studies. The source code and user documentation are available at <u>https://github.com/MDU-PHL/ngmaster</u>.

## DATA SUMMARY

1. The Python source code for *NGMASTER* is available from GitHub under GNU GPL v2. (URL: <u>https://github.com/MDU-PHL/ngmaster</u>)
2. The software is installable via the Python “pip” package management system. Install using “pip install --user git+https://github.com/MDU-PHL/ngmaster.git”
3. Sequencing data used are available for download from the EBI European Nucleotide Archive under BioProject accessions <u>PRJEB2999</u>, <u>PRJNA29335</u>, <u>PRJNA266539</u>, <u>PRJNA298332</u>, and <u>PRJEB14168</u>.

## IMPACT STATEMENT

Whole-genome sequencing (WGS) offers the potential for high-resolution comparative analyses of microbial pathogens. However, there remains a need for backward compatibility with previous molecular typing methods to place genomic studies in context. NG-MAST is currently the most widely used method for epidemiologic surveillance of *Neisseria gonorrhoeae*. We present *NGMASTER*, a command-line software tool for performing MultiAntigen Sequence Typing (NG-MAST) of *Neisseria gonorrhoeae* from WGS data. This tool is targeted at clinical and research microbiology laboratories that have performed WGS of *N. gonorrhoeae* isolates and wish to understand the molecular context of their data in comparison to previously published epidemiological studies. As WGS becomes more routinely performed, *NGMASTER* was developed to completely replace PCR-based NG-MAST, reducing time and labour costs.

## INTRODUCTION

*Neisseria gonorrhoeae* is one of the most common sexually transmitted bacterial infections worldwide. There is growing concern about the global spread of resistant epidemic clones, with extensively drug-resistant gonorrhoea being listed as an urgent antimicrobial resistance threat (CDC, 2013; WHO, 2014).

Multi-Antigen Sequence Typing of *N. gonorrhoeae* (NG-MAST) has been important in tracking these resistant clones, such as the NG-MAST 1407 clone associated with decreased susceptibility to third-generation cephalosporins (Unemo & Dillon, 2011). It involves sequence-based typing using established PCR primers of two highly variable and polymorphic outer membrane protein genes,*porB* and *tbpB* by comparing the sequences to an open-access database (http://www.ng-mast.net/) (Martin *et al.*, 2004). Although NG-MAST is the most frequently performed molecular typing method for *N. gonorrhoeae*, it requires multiple PCR amplification and sequencing reactions, making it more laborious than other typing methods (Heymans *et al.*, 2012).

Whole-genome sequencing (WGS) is increasingly being used for molecular typing and epidemiologic investigation of microbial pathogens as it provides considerably higher resolution. A number of studies using genomic data to understand the epidemiology of *N. gonorrhoeae* have already been published (Grad *et al.*, 2014) (Demczuk *et al.*, 2015) (Ezewudo *et al.*, 2015) (Demczuk *et al.*, 2016). However, the ability to perform retrospective comparisons with previous epidemiological studies is reliant on conducting both traditional typing (such as NG-MAST) as well as more modern WGS analyses on the same isolates.

*NGMASTER* is a command-line software tool for rapidly determining NG-MAST types *in silico* from genome assemblies of *N. gonorrhoeae*.

## DESCRIPTION

*NGMASTER* is an open source tool written in Python and released under a GPLv2 Licence. The source code can be downloaded from Github (<u>https://github.com/MDU-PHL/ngmaster</u>). It has two software dependencies: *isPcr* (<u>http://hgwdev.cse.ucsc.edu/~kent/src/</u>) and BioPython (Cock *et al.*, 2009), and uses the allele databases publicly available at <u>http://www.ng-mast.net/</u>, which *NGMASTER* can automatically download and update locally for running.

*NGMASTER* is based on the laboratory method published by Martin *et al.* (Martin *et al.*, 2004), and uses *isPcr* to retrieve allele sequences from a user-specified genome assembly in FASTA format by locating the flanking primers. These allele sequences are trimmed to a set length from starting key motifs in conserved gene regions, and then checked against the allele databases. Results are printed in machine readable tab-or comma-separated format.

## METHODS AND RESULTS

*NGMASTER* was validated against 630 publicly available *N. gonorrhoeae* genome sequences derived from published studies (Table 1). This included 8 well characterised WHO reference genomes with published data and 50 local isolates that had undergone “traditional” NG-MAST by PCR and Sanger sequencing (Martin *et al.*, 2004). A further 572 isolates that had undergone manual *in silico* NG-MAST from WGS data (Demczuk *et al.*, 2015; Demczuk *et al.*, 2016; Grad *et al.*, 2014), including the fully assembled reference genome NCCP11945, were also tested. Raw WGS data for these sequences were retrieved from the European Nucleotide Archive (ENA). Average sequencing depth was >30x for all ENA sequences, with a combination of 100 bp, 250 bp and 300 bp paired-end Illumina reads. Local isolates also underwent WGS on the Illumina MiSeq/NextSeq using Nextera libraries and manufacturer protocols, with an average sequencing depth >50x. The raw sequencing reads for these local isolates have been uploaded to the ENA (BioProject accession PRJEB14168).

**Table 1:**
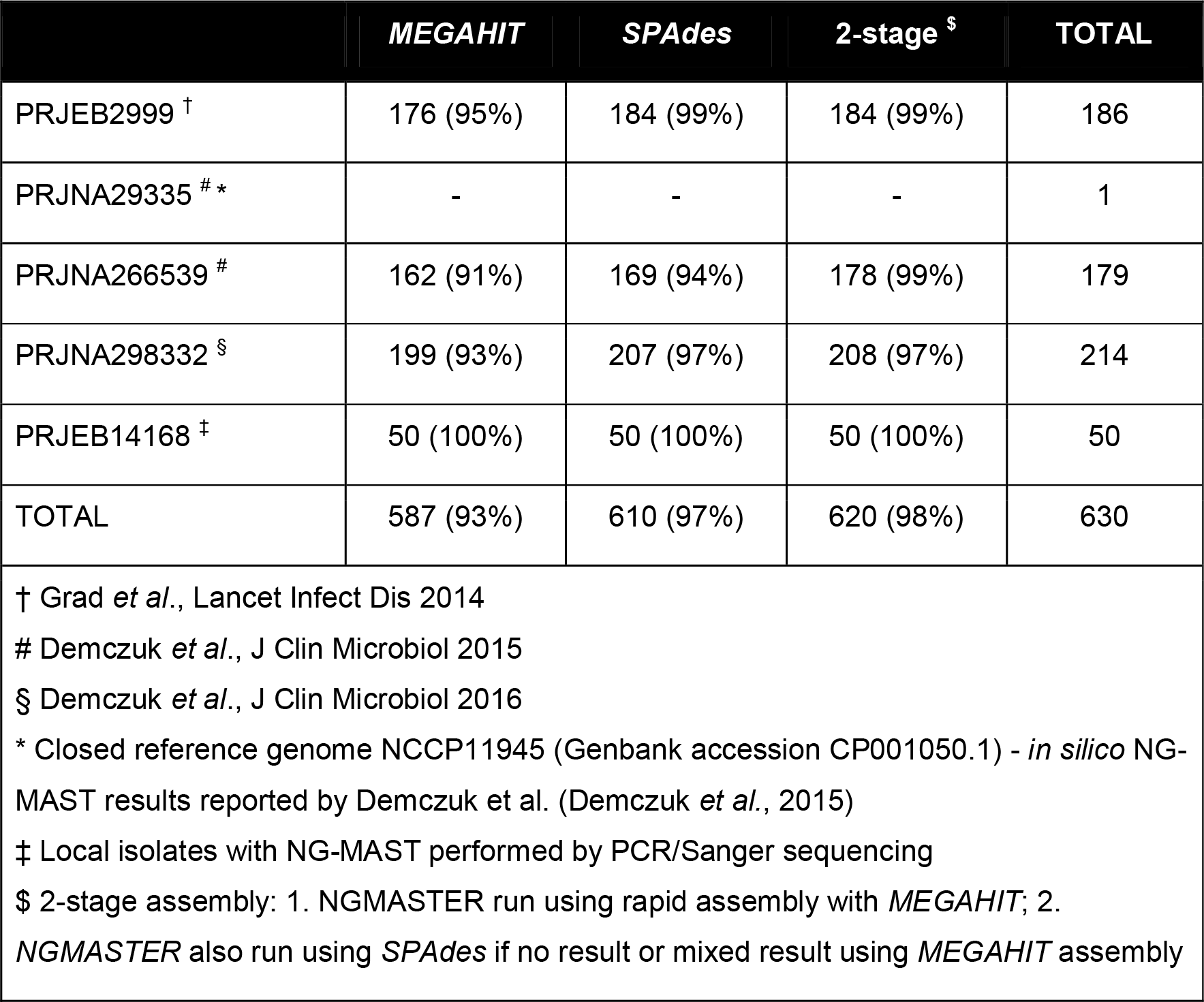
Concordance between *NGMASTER* results from draft genome assemblies using *MEGAHIT* and *SPAdes*, and previously published NG-MAST results.

Sequencing reads were trimmed to clip Illumina adapters and low-quality sequence (minimum Q20) using *Trimmomatic* v0.35 (Bolger *et al.*, 2014). Draft genomes were assembled *de novo* with *MEGAHIT* v1.0.3 and *SPAdes* v3.7.1 (Li *et al.*, 2015) (Bankevich *et al.*, 2012) to investigate whether the faster, but approximate genome assembler, *MEGAHIT*, would be sufficient for *NGMASTER*. A list of the commands and parameters used is included in the Appendix 1.

The *de novo* assembled draft genomes and the fully assembled NCCP11945 reference genome in FASTA format were used as input to *NGMASTER* with the overall results shown in Table 1. Complete *NGMASTER* results with sequencing and assembly metrics are included in Appendix 2. Running *NGMASTER* on 630 genome assemblies using a single Intel(R) Xeon(R) 2.3GHz CPU core completed in less than two minutes.

Overall, *NGMASTER* assigned NG-MAST types that were concordant with published results for 93-97% of the tested *N. gonorrhoeae* genomes using *MEGAHIT* or *SPAdes* assemblies. Notably, comparisons with results from traditional NG-MAST were 100% concordant (57/57). Reasons for discordant results are shown in Table 2. In general, using *SPAdes* assemblies with *NGMASTER* resolved more NG-MAST types than when using *MEGAHIT* assemblies. However, 10 genomes assembled with *SPAdes* were found to have assembly errors in either *por* or *tbpB* introduced in the repeat resolution stage, resulting in discordant NG-MAST types for those isolates (major errors). Running *NGMASTER* on preliminary *SPAdes*-assembled contigs prior to this process (“before_rr.fasta”) alleviated these major errors, and were concordant with the *MEGAHIT* results and published data (Appendix 2). In contrast, minor errors (due to incomplete NG-MAST types or multiple alleles detected) were more frequent using *MEGAHIT* assemblies, particularly those with poor assembly metrics (e.g. >500 contigs, N50 <10 kbp). When *MEGAHIT* assemblies successfully produced complete *NGMASTER* results, these NG-MAST types were highly concordant with the published results.

**Table 2:** Reasons for discordant results between *NGMASTER* and published data using *SPAdes* assemblies

| Reason for discordant result | *MEGAHIT* | *SPAdes* |
| --- | --- | --- |
| ***Major errors (incorrect result)*** |  |  |
| Assembly error | 0 | 10 |
| ***Minor errors (incomplete/missing result)*** |  |  |
| Alternate conserved key motif | 1 | 1 |
| Multiple alleles detected | 6 | 2 |
| Allele not detected | 29 | 0 |
| ***Errors in published data*** |  |  |
| Possible sequence mix-up in published data | 4 | 4 |
| Probable transcription error in published data | 1 | 1 |
| Error in published data | 1 | 1 |

To overcome this issue, a two-stage assembly approach was also tested, where a draft genome was first assembled using *MEGAHIT* for initial testing. If a complete NG-MAST result was obtained, this was recorded as the final result for that isolate. If the result was incomplete or suggested multiple alleles were present, the genome was also assembled using *SPAdes*. Using this combined approach, 620/630 (98%) NG-MAST types derived from *NGMASTER* were concordant with the published results, with only 42 genomes requiring additional assembly with the slower, but more thorough *SPAdes* assembler.

For the remaining 10 discordant results, seven of these were likely due to errors in the published data, including for NCCP11945. A further two isolates were found to have multiple *tbpB* alleles in both *SPAdes* and *MEGAHIT* assemblies, with the dominant allele (indicated by higher read coverage and better flanking assembly) matching the published result. The *tbpB* allele for the final isolate was not able to be determined by *NGMASTER* due to a mutation in the conserved starting key motif required for sequence trimming to a standard size.

## ISSUES WITH IMPLEMENTATION

The NG-MAST procedure involves sequencing the internal regions of *por* and *tbpB* that encode two variable outer membrane proteins. The sequences are trimmed to a standard length from a starting key motif in conserved regions of each gene. However, despite being relatively conserved, a number of variations of this starting motif appear in the NG-MAST database (Fig. 1), causing one discordant result (Table 2). Some sequences appeared to lack a *tbpB* gene due to the presence of non-typeable *tbpB* genes acquired from *N. meningitidis*, though this was also noted in the published data. Another source of discordant results were genomes that appeared to have multiple alleles, suggesting isolate contamination or polyclonal infection.

**Figure 1:**
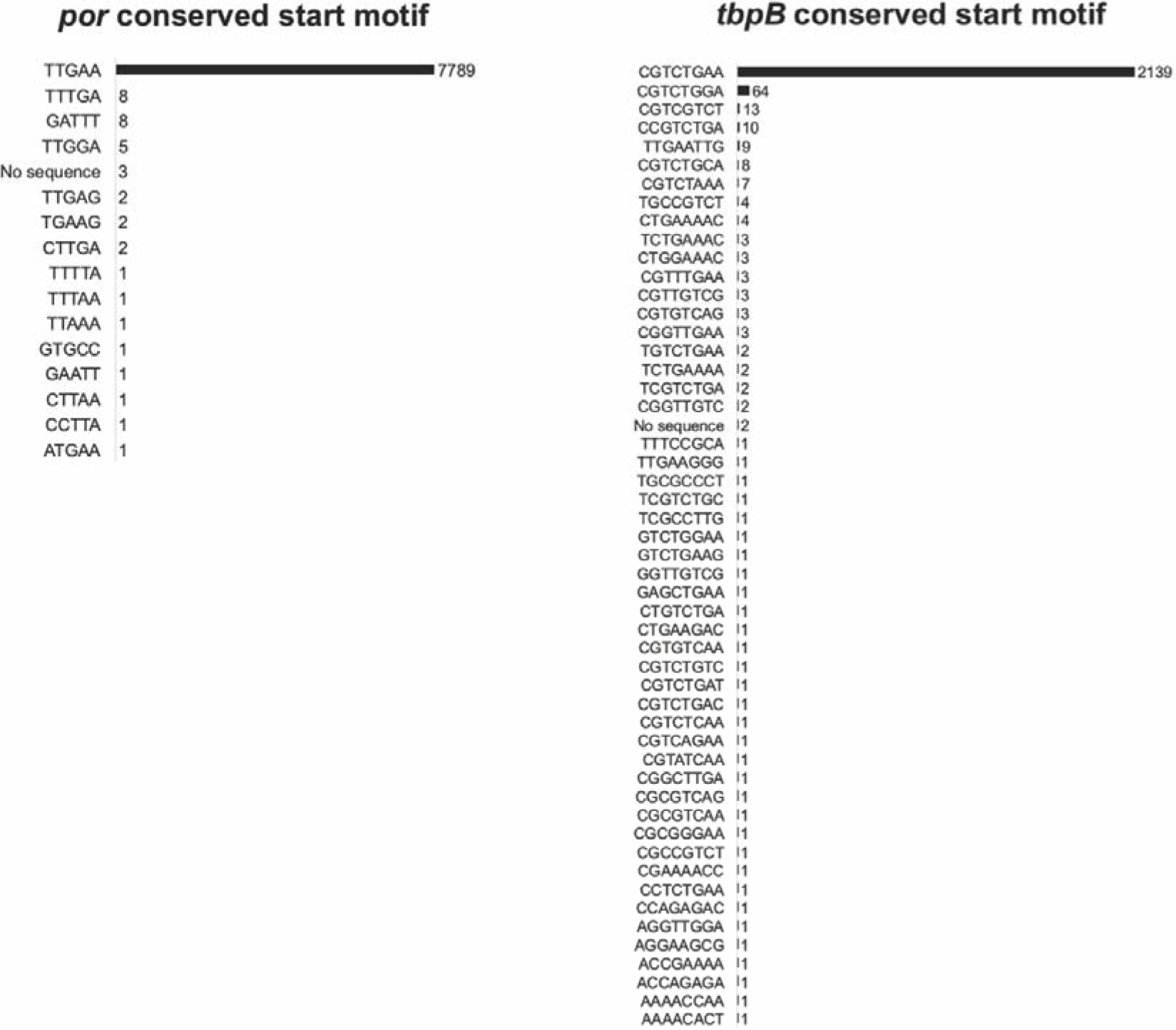
Number and frequency of alternate starting key motifs within “conserved” gene regions for trimming allele sequences.

A number of isolates were found to have novel alleles or allele combinations that were not in the most recent version of the database available at <u>http://www.ng-mast.net</u>. For convenience, *NGMASTER* includes an option to save these allele sequences in FASTA format for manual submission to the database and allele type assignment.

Notably, results were dependent on the accuracy and quality of the *de novo* draft genome assembly. It should be noted that for this study, draft genomes were assembled *de novo* using relatively standard parameters for *MEGAHIT* and *SPAdes* without post-assembly error checking (see Appendix 1). We were alerted to the presence of *SPAdes* assembly errors after finding the corresponding *MEGAHIT* assemblies produced different *NGMASTER* results. Concordant results were able to be obtained for each of these genomes after identifying and correcting assembly errors through re-mapping each isolate’s reads back to the respective draft *SPAdes* assembly. Results from running *NGMASTER* on the *SPAdes* interim “before_rr.fasta” contigs also produced concordant results. Assuming accurate closed genome assemblies are used with an accurate and well curated database, based on our testing, we anticipate that *NGMASTER* would produce NG-MAST results that were >99% if not 100% accurate.

## CONCLUSION

*NGMASTER* rapidly and accurately performs *in silico* NG-MAST typing of *N. gonorrhoeae* from assembled WGS data, and may be a useful command-line tool to help contextualise genomic epidemiological studies of *N. gonorrhoeae*.

## Acknowledgements

We wish to acknowledge the contributions of Helen Heffernan at the Institute of Environmental Science and Research, New Zealand, and Kerrie Stevens and the Molecular Diagnostics section staff at the Microbiological Diagnostic Unit Public Health Laboratory, Australia who assisted in providing NG-MAST data.

